# Low photosynthetic rate under low light stress inhibited sucrose distribution and transportation to grain

**DOI:** 10.1101/2022.08.02.502494

**Authors:** Zhichao Sun, Wenjie Geng, Baizhao Ren, Bin Zhao, Peng Liu, Jiwang Zhang

## Abstract

Under the condition of low light, the yield of summer maize decreased significantly, but the decrease of yield under low light stress was not only caused by the lack of photosynthetic assimilates in leaves, but also the transportation and utilization of assimilates by stems and grains. In this study, we investigated the effects of low light stress on leaves, stems and grains of summer maize and the relationship between them. The results showed that the synthesis ability of sucrose and export sucrose to grains ability in leaves decreased under low light. Due to dry matter transfer, the number and area of small vascular bundles in spike node and shank decreased, which restricted the translocation of photoassimilates to grains at filling stage. The activities of SUS and AGPase was decreased in grains under low light stress, which limited the availability of sucrose. The process of leaf synthesis, sucrose loading and sucrose utilization in grains was affected under low light, resulting in relatively higher sucrose concentration in grains than in leaves, forming a “leaf low” - “grain high” sugar concentration gradient, resulting in the opposite hydrostatic pressure, and then feedback inhibition of sucrose output in leaves, reducing sucrose loading and transportation rate.

**Hightlight:** The key factor of low light stress reducing summer maize yield was the decrease of leaf photosynthetic rate, resulting in insufficient grain dry matter supply. The sugar concentration gradient between leaves and grains further restricted the sucrose transport from leaves to grains.

## Introduction

It was expected that by 2050, the global population would reach 9 billion (Godfray *et al*., 2010) and the Novel Coronavirus pandemic (Lamichhane and Reay-Jones, 2021) would pose a severe challenge to global food supply and food security. As one of the most important crops in the world, maize is an important source of food and feed for human beings. However, in recent years, environmental pollution has increased and extreme weather has occurred frequently, resulting in a significant reduction in the total solar radiation and effective sunshine hours in China (Ramanathan and Feng, 2009; Shao *et al*., 2021; Yang *et al*., 2021). Insufficient light during the growth period of summer maize resulted in a significant decrease in yield (Zhang *et al*., 2006). Therefore, it is necessary to clarify the mechanism of low light affecting maize yield formation and provide coping strategies for future climate change.

The main source of grain yield was photoassimilates formed after silking (Shi *et al*., 2015). The expression of PEPC related genes was susceptible to light intensity (Chollet *et al*., 1996), and the activation of Rubisco was limited by light-dependent adenosine triphosphate (ATP) (Wu *et al*., 2014). Under low light conditions, the activities of photosynthetic enzymes such as phosphoenolpyruvate carboxylase (PEPC), ribulose bisphosphate carboxylase oxygenase (Rubisco) and NADP-dependent malic enzyme (NADP-ME) was decreased significantly (Zhang *et al*., 2007 ; Sharwood *et al*., 2014), photoassimilates accumulation decreased. The triose phosphate produced during the Calvin cycle has three places, one was involved in the regeneration of ribulose diphosphate, the second was the synthesis of starch in the chloroplast by adenosine diphosphate glucose pyrophosphorylase (AGPase) and other enzymes, and the third was transported to the cytoplasm through the triose phosphate translocator (TPT), sucrose phosphate synthase (SPS) and other enzymes to synthesize sucrose and then transported to the grain. The output rate of triose phosphate to the cytoplasm determines the synthesis and output rate of sucrose, but it is difficult to determine the output rate due to the limited technology at present. Therefore, the activity of enzymes was used to determine the location of triose phosphate. The changes of enzyme activities such as AGPase and SPS are still unknown under low light conditions. Grain filling substances mainly came from two aspects, one was the transformation of storage substances in organs such as stems, and the other was the accumulation of photoassimilate after flowering, of which the second aspect accounted for 70-90%. Lack of light led to insufficient production of dry matter, and grains were required to transport more nutrients from vegetative organs. The transport of assimilates in vegetative organs before silking increased (Wang *et al*., 2020; Yang *et al*., 2021). However, the effect of increased dry matter transport in stem on long-distance transport of photoassimilates after flowering is unknown.

The transportation of carbohydrates in plants from “source” tissue to “sink” tissue mainly included three stages. The loading of phloem in “source” tissue was the starting point of long-distance transportation of nutrients (Zhang *et al*., 2015). Previous studies have shown that maize leaves use apoplast transport based on ultrastructural, physiological and genetic data (Bezrutczyk *et al*., 2021). Sucrose was first synthesized in the cytoplasm of mesophyll cells (MS) and moved from MS to phloem parenchyma cells (PP) via plasmodesmata (Braun, 2022). It was then excreted by sucrose transporters (ZmSWEET13a, ZmSWEET13b, and ZmSWEET13c) (Bezrutczyk *et al*., 2018) and entered the companion cell (CC)-sieve element (SE) complex (Slewinski *et al*., 2009) via SUT1. Then it was transported to grains in SE (Braun, 2022). Previous studies found that stress changed the photoassimilate transport rate by altering the phloem loading process. After low temperature stress, the expression level of maize genes involved in plasmodesmata operation changed significantly (Bilska-Kos *et al*., 2016). Low light inhibited the growth of SE and CC in nectarine phloem tissue, and the density of plasmodesmata decreased or even did not exist (Wang and Huang, 2003). The expression of SUT1 in different parts was changed under different degrees of low light stress (Ishibashi *et al*., 2014). At present, there are few reports about the effect of low light on phloem loading of summer maize leaves.

The pressure flow model has been widely accepted to explain sucrose transport dynamics (Patrick, 2013). After sucrose accumulates to a high concentration in SEs of leaf, water flows into the SEs from xylem (X) through infiltration to produce relatively high hydrostatic pressure. In grains, sucrose was unloaded from the phloem, resulting in the diffusion of water into surrounding cells and subsequent reduction of hydrostatic pressure. Under low light conditions, the photosynthetic rate of leaves decreased. What sugar concentration gradient will result from the competition between leaves and grains for limited sucrose? Does the difference in sugar concentration between leaves and grains affect phloem transport dynamics? These are questions for further exploration. Studies by Chen et al. (2011) showed that *AtSWEET13* guaranteed the high efflux rate of Arabidopsis at low photoassimilation rate under low light conditions. Does summer maize have an adaptive mechanism to ensure the balance of sucrose efflux and absorption between PP and CC-SE under low light?

Therefore, this experiment simulated low light stress by artificial shading to clarify (1) how maize coordinates the distribution of Calvin cycle products between leaves and grains; (2) the effects of low light on loading process of sucrose phloem; (3) the effects of sucrose concentration gradient formed between leaves and grains on sucrose output in leaves; (4) Whether the pre-flowering dry matter transport increased and the effect of increased transport on long-distance sucrose transport in the late stage; (5) the performance of summer maize to adapt to low light stress, to provide theoretical basis for the stress-resistant and high-yield cultivation technology of summer maize.

## Materials and methods

### Experimental procedures

The experiment was conducted at the State Key Laboratory of Crop Biology and the Experimental Farm of Shandong Agricultural University (36°17’N, 117°17’E) in 2020-2021. The soil type was brown loam, and the basic fertility of 0-20 cm soil before sowing was as follows: soil organic matter 9.2 g kg^-1^, total nitrogen 0.7 g kg^-1^, total phosphorus 0.8 g kg^-1^, available nitrogen 78.7 mg kg^-1^, available phosphorus 35.6mg kg^-1^, available potassium 84.5 mg kg^-1^., Summer maize hybrid Denghai 605(DH605) was selected as experimental material, and the planting density was 67, 500 ha^-1^ plants. Two experimental treatments, S (shading from flowering to maturity stage) and CK (natural light control), were set under field conditions, and shading was 60% (simulated cloudy day), which was achieved by scaffolding and black nylon net (Jia Wan Ying Trading Co., Ltd, Linyi City, China) with light transmission rate of 40%. A distance of 2 m was kept between the shading net and the maize canopy to ensure that the field microclimate in the shading shed was basically consistent with the control conditions. The plot area was 27 m^2^ (9 m×3 m) and 3 replicates were set. Sowing on June 8th, irrigation method was sprinkler irrigation, spraying acetochlor (20 %) and atrazine (20 %) to control weeds before emergence, spraying difenoconazole (10 %) to control rust and brown spot. At the 6-leaf stage (V6), 750 kg ha^-1^ compound fertilizer (N: P_2_O_5_: K_2_O= 28:6:6) was applied.

### Field microclimate

At tasseling stage (VT), the light intensity at the top, ear and bottom of maize was measured by digital illuminometer (TES-1332A, TES Co. Ltd, Taiwan, China), the CO_2_ concentration at ear was measured by CIRAS-III (PP System, Hansatech, USA), temperature and humidity recorder (GSP-6, Elitech Co. Ltd, Jiangsu, China) measured the temperature and humidity of ear, used soil temperature and humidity recorder (SY-HWS, Yashi, Co. Ltd, Hebei, China) to measure the temperature and humidity of soil, and used portable weather instrument (NK-5500, KESTREL Instruments Co. Ltd, USA) to measure the wind speed of ear. Five replicates were measured for each treatment, and the above indicators were measured at 11:00 (Table 1) (Gao *et al*., 2017).

**Table 1.** Effects of low-light stress on the microclimate in experimental field. Different lowercase letters in the same column indicate significant difference at P < 0. 05 by LSD test.

| Treatment | Light intensity ($\mu\text{mol m}^{-2} \text{s}^{-1}$) | | | Air speed<br>($\text{m s}^{-1}$) | Temperature<br>($^{\circ}\text{C}$) | Relative<br>humidity (%) | CO<sub>2</sub> concentration<br>($\mu\text{mol mol}^{-1}$) |
| --- | --- | --- | --- | --- | --- | --- | --- |
|  | Canopy | Ear | Ground |  |  |  |  |
| CK | 799.2a | 284.a | 121.0a | 0.4a | 32.4a | 75.4a | 284.9a |
| S | 316.1b | 137.2b | 57.6b | 0.3a | 31.1a | 77.0a | 298.6a |

S: Shading from flowering to maturity stage; CK: Normal light from flowering to maturity stage. Different lowercase letters in the same column indicate significant difference at $P < 0.05$ by LSD test.

### Yield

Thirty ears were harvested continuously in 3 rows in the middle of each plot and used to measure yield after natural drying (converted to 14% water content) (Shi *et al*., 2015). The number of harvested ears was the number of effective ears for field investigation. The formula for calculating yield was: Yield (kg ha^-1^) = number of ears (ha^-1^) × number of grains per ear × 1000-grain weight (g) /10^6^× (1-water content %) / (1-14%)

### Dry matter accumulation and distribution

At VT stage and physiological maturity stage (R6), 5 representative plants with the same growth status were selected from each plot and divided into stems, leaves (VT) or stems, leaves and ears (R6). The plants were placed in the oven (DHG-9420A, Bilon Instruments Co. Ltd, Shanghai, China) at 110 °C for degreening and dried at 80 °C until constant weight weighing. Dry matter weight in stem (DW_S_), dry matter weight in leaf (DW_L_), dry matter weight in grain (DW_G_), proportion of dry matter in stem (DWP_S_), proportion of dry matter in leaf (DWP_L_), proportion of dry matter in grain (DWP_G_) and other indicators representing the accumulation and distribution of dry matter were calculated by the following formula (Yang *et al*., 2021).

Dry matter proportion of each organ (%) = dry weight of vegetative organ (g plant^−1^) / total dry weight of plant (g plant^−1^) × 100;

Dry matter translocation of pre-flowering (g plant^−1^) (TBA) = dry matter accumulation at flowering (g plant^−1^) –dry matter accumulation of vegetative organs at maturity (g plant^−1^);

Dry matter transport rate of pre-flowering (%) (TRBA) = TBA (g plant^−1^) / dry matter accumulation in shoot at flowering (g plant^−1^) × 100;

Contribution rate of pre-flowering dry matter to grain (%) (TFGR) = TBA (g plant^−1^) / kernel dry matter weight at maturity (g plant^−1^) × 100;

The amount of assimilated dry matter after flowering (g plant^−1^) (PAA) = dry matter accumulation in shoot at maturity (g plant^−1^) – dry matter accumulation in shoot at flowering (g plant^−1^) = kernel dry matter weight at maturity (g plant^−1^) – TBA (g plant^−1^);

Contribution rate of dry matter assimilation to grain after flowering (%) (PAGR) = PAA (g plant^−1^) / kernel dry matter weight at maturity (g plant^−1^) × 100.

### Gas exchange measurements

At R3 stage, 10 representative plants with the same growth status were selected for each treatment, and gas exchange parameters such as net photosynthetic rate (Pn), stomatal conductance (Gs), intercellular carbon dioxide concentration (Ci) and transpiration rate (E) were measured by CIRAS-III from 9: 00 to 11: 00 (Huang *et al*., 2020).

### Observation of Kranz anatomy and determination of plasmodesmata densities

At R3 stage, the ear leaves of 3 representative plants with consistent growth were selected. A square leaf (0.5cm × 0.5cm) was taken from the middle of the leaf (avoiding veins) and fixed with 2.5% glutaraldehyde fixative solution. Air was pumped until the cut pieces settled and fixed at 4 °C for 24 h. The material was washed with PBS buffer for 5 times, 20 min each, and then transferred to O_S_O_4_ for 4.5h fixation. The leaf tissues were then washed with PBS buffer for 5 times, dehydrated with conventional gradient ethanol, replaced with epoxy propane and embedded with resin, and polymerized for 3 days at different temperatures. After treatment, the leaf tissues were sectionalized with LKB-5 ultrafine microtome, stained with uranyl acetate and lead citrate, and observed under transmission electron microscope (JEM-1400Plus, JEOL Co, Japan). The density of plasmodesmata was determined according to the number of plasmodesmata per 5 μm (including MS-BS, BS-BS, BS-PP) (Chen *et al*., 2020). After the embedded samples were treated with saw blade and blade, the semi-thin sections (thickness 2 μm) were cut by an automatic semi-thin rotary cutting machine (LEICA RM2265, Leica Microsystems Co., Ltd., Germany) and triangular glass knife. A drop of double-distilled water was dropped on the clean slide in advance, and the sliced semi-thin slices were placed on the double-distilled water with tweezers. The slices were placed on a digital hotplate (Benchmark H3760-H, Fotronic Co, USA), and dried at 50 °C. Then the slices were further microscopically examined using a fluorescence microscopy imaging system (OlympusBX51, Olympus Co., Tokyo, Japan) and photographed to calculate the number and area of the Kranz anatomy with Image J (Version 1.8.0, National Institutes of Health) software (Ren *et al*., 2016).

### RNA isolation, reverse transcription, and real time PCR (qPCR) analysis

At R3 stage, 3 representative plants of each treatment were selected, and the ear-position leaves were stored at -80°C for the determination of relative gene expression levels of *SUT1, SWEET13a, SWEET13b* and *SWEET13c*. Total RNA was extracted using an HiPure Plant RNA Mini Kit (MGBio). RNA concentration and purity were assessed using NanoDrop2000 microultraviolet spectrophotometer (Thermo Scientific, USA). Afterwards, the isolated RNA sample was used as template for complementary DNA (cDNA) synthesis using oligo (dTs) and abm’s proprietary OneScript^®^ Hot Reverse Transcriptase (Invitrogen). For qPCR, Bestar^®^ SybrGreen qPCR Mastermix (DBI^®^ Bioscience, Germany) was used in the reaction mixture according to the manufacturer’s instructions, and reactions were performed in eight-link boards on QuantStudio™ 6 Flex Real-Time PCR System (Applied Biosystems, USA) with cycling conditions (50 °C 2 min, 95 °C 2 min, 95 °C 10 s, 60 °C 20 s, 40 cycles, 95 °C 15 s, 55 °C 1 min). Primers were listed in Supplementary Table S1 at JXB online. 3 biological replicates were determined for each sample, and 3 technical replicates were performed for each biological replicate. The obtained data was used to calculate the relative expression of genes by 2^−ΔΔCt^ method (Livak and Schmittgen, 2001).

### Stalk trait measurements

At R3 stage, 4 representative plants were selected for each treatment and about 1.5 cm stem at the middle part of the spike node and shank was fixed in the Carnoy fixative and stored with 70% ethanol. Thin slices were cut using the method of free hand section and stained with safranin. The slices were further microscopically examined using a fluorescence microscopy imaging system (OlympusBX51, Olympus Co., Tokyo, Japan) and photographed to calculate the number and area of big, small vascular bundle per visual field with Image J (Version 1.8.0, National Institutes of Health) software (Guo *et al*., 2016).

### Grain filling process measurement

A total of 30 plants with similar growth patterns that silked on the same day were labeled. 5 tagged ears from each plot were sampled at 10-day intervals from silking until maturity stage; 100 grains were sampled from the middle part of the ear and oven-dried at 85°C to a constant weight. Grain filling process was simulated by Logistics equation: W = a / [1 + b × exp (–c × t)]. where W = grain weight (g); t = number of days after silking; a was the potential kernel weight (g), b and c were coefficients determined by regression. The grouting parameters were calculated by the following formula (Shi *et al*., 2013):

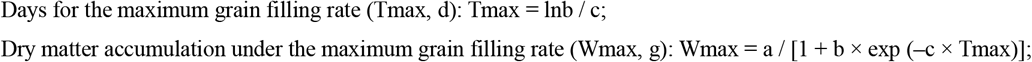

Maximum grain filling rate (Gmax, g 100kernel^−1^ d^−1^):

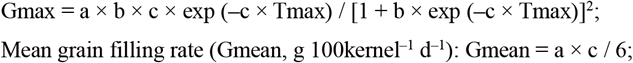

Active grain filling period (D, d): D = a / Gmean.

### Assay of sucrose, and starch contents

At R3 stage, 3 plants were sampled from each treatment and separated into ear leaves and ears. All samples were dried, weighed, and ball milled. 100 mg tissue samples were extracted directly in 7 mL boiling water for 20 min, the supernatant was collected, and the residues were extracted a second time in 7 mL boiling water for 20 min. The extract was then diluted to 50 mL constant volume with deionized water and named solution A. In addition, 4 mL water and 2 mL 9.2 N HClO_4_ were added to the residues, then were placed in boiling bath for 20 min. After this, the supernatant was collected, and the residues were extracted a second time in 5 mL water and 1 mL 9.2N HClO_4_ in boiling bath for 20 min. The extract was then diluted to 50 mL constant volume with deionized water and named solution B. For sucrose content analysis, 100 μL solution A was reacted with 100 μL KOH (30%) for 10 min in a boiling water bath, cooled, and added to 3 mL anthracenone solution after cooling, and absorbance at 620 nm was measured. For starch content analysis, 100 μL solution B was reacted with 3 mL anthracenone solution for 10 min in a boiling water bath and cooled, and absorbance at 620 nm was measured (Hu *et al*., 2022).

### Enzyme activity assays

3 representative plants were selected in each treatment at R3 stage, and fresh samples of ear leaves (avoiding leaf veins) and grains were taken respectively, frozen with liquid nitrogen and stored in − 80 °C refrigerator. PEPC, NADP-ME, Rubisco, SPS activities in leaves and fructose content, sucrose synthase (SUS), cell wall invertase (CWI) activities in grains were operated according to manufacturer instructions (Cominbio, www.cominbio.com).

### Data analysis

Statistical analyses were performed in Excel 2019 (Microsoft, Redmond, WA, United States) and IBM SPSS Statistics 21.0 (IBM Corporation, Armonk, NY, USA). One-way ANOVA with the least significant difference test (LSD, α = 0.05) was used to test the difference of yield, grain weight and maximum filling rate among different treatments. Figures were produced with Sigmaplot 14.0 (Systat Software, San Jose, CA).

## Results

### Yield, dry matter accumulation and distribution

The yield of summer maize was significantly reduced under low light stress, and the trend was consistent in the two years. Compared with CK, harvest ear number, grain number per ear, 1000-grain weight and yield of S treatment were decreased by 0.9 %, 41.3 %, 13.3 % and 49.28 % on average (Table 2). The ear picture of the two treatments was shown in Figure 1. Dry matter accumulation and grain distribution ratio of summer maize were significantly reduced under low light stress (Figure 2). The trend was consistent in the two years. The rust was serious in 2020, and the data in 2021 were analyzed. At VT stage, DWP_S_ and DWP_L_ in CK treatment were 63.8 % and 36.2 %. At R3 stage, DW_S_ DW_L_ and DW_G_ of S treatment were decreased by 16.4 %, 18.2 % and 59.8 % compared with CK, and DWP_S_, DWP_L_ and DWP_G_ were decreased by-10.5 %, -4.8 % and 12.9 %. At R6 stage, DW_S_, DW_L_ and DW_G_ of S treatment decreased by 18.8 %, 14.2 % and 51.4 %, DWP_S_, DWP_L_ and DWP_G_ decreased by -7.1 %, -4.9 % and 12.6 %. Dry matter production was decreased significantly after flowering under low light stress, and dry matter in grain mainly came from dry matter accumulated before flowering. TBA, TRBA, TFGR, PAA and PFGR of S treatment were significantly lower than those of CK by -98.0%, -15.6%, -47.1%, 79.5% and 47.1%, respectively (Figure 2A, B).

**Table 2.**
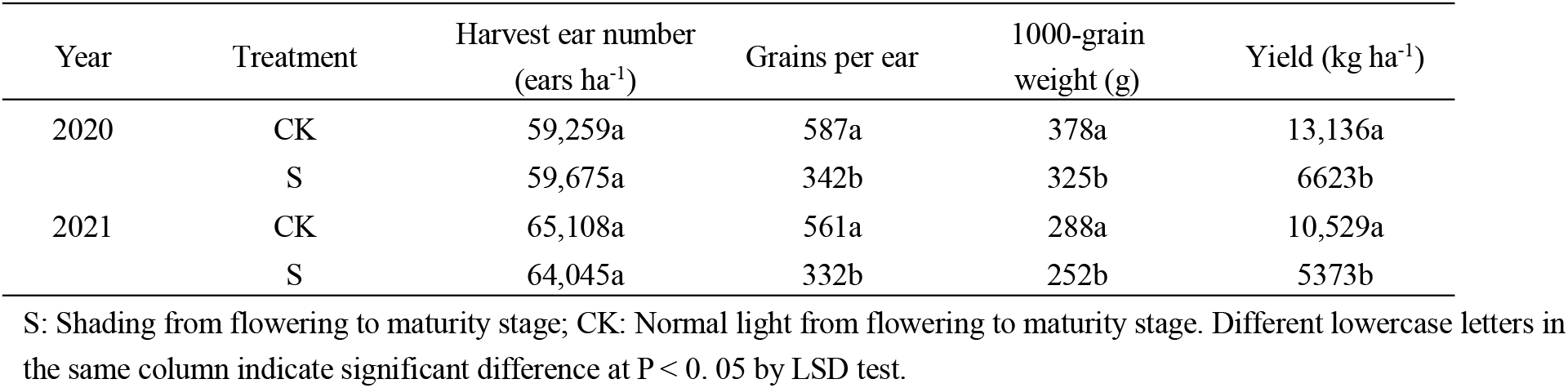
Effects of low-light stress on yield and its components of summer maize. Different lowercase letters in the same column indicate significant difference at P < 0. 05 by LSD test.

| Year | Treatment | Harvest ear number<br>(ears $\text{ha}^{-1}$) | Grains per ear | 1000-grain<br>weight (g) | Yield ($\text{kg ha}^{-1}$) |
| --- | --- | --- | --- | --- | --- |
| 2020 | CK | 59,259a | 587a | 378a | 13,136a |
|  | S | 59,675a | 342b | 325b | 6623b |
| 2021 | CK | 65,108a | 561a | 288a | 10,529a |
|  | S | 64,045a | 332b | 252b | 5373b |

**Figure 1.**
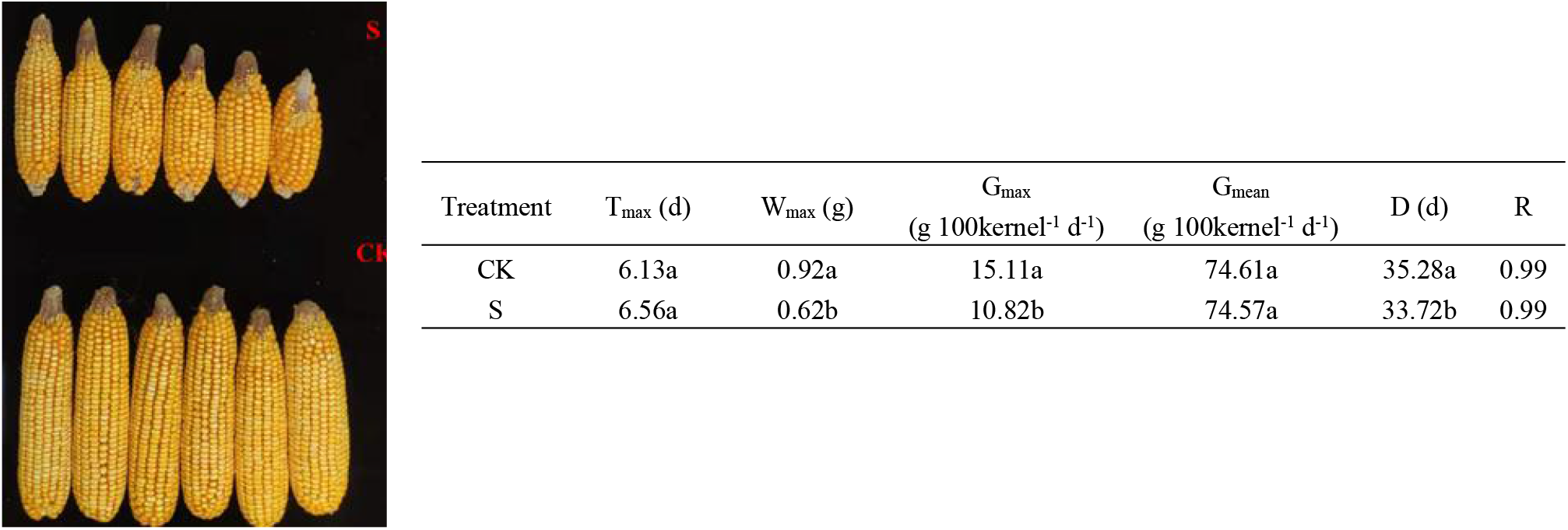
Effects of low-light stress on the characteristic parameters of grain-filling of summer maize. S: Shading from flowering to maturity stage; CK: Normal light from flowering to maturity stage; T_max_: Days for the maximum grain filling rate; W_max_: Dry matter accumulation under the maximum grain filling rate; G_max_: Maximum grain filling rate; G_mean_: Mean grain filling rate; D: Active grain filling period. Different lowercase letters in the same column indicate significant difference at P < 0. 05 by LSD test.

**Figure 2.**
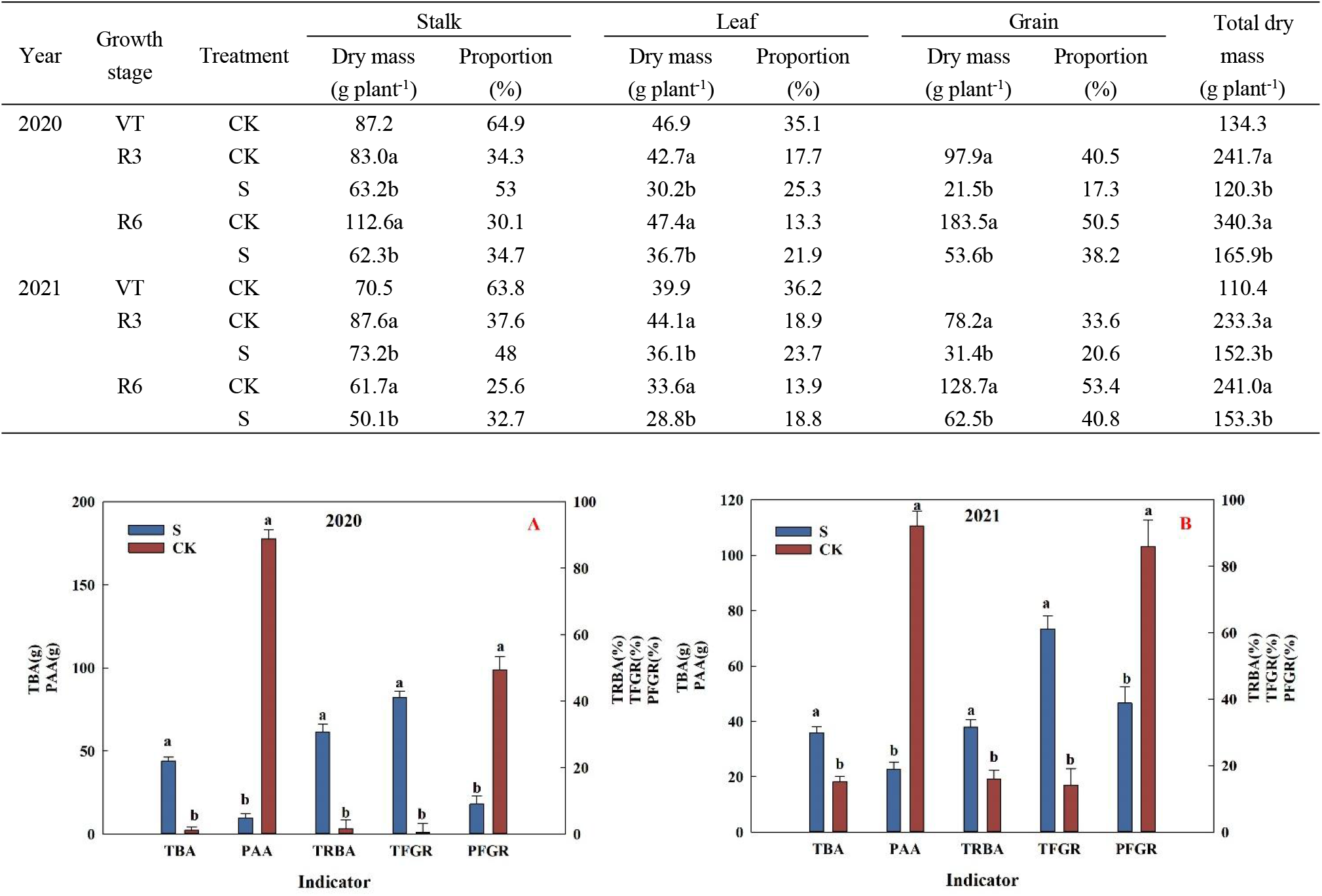
Effects of low-light stress on dry matter accumulation and distribution of summer maize. (A) Effects of low light stress on dry matter accumulation and distribution of summer maize in 2020. (B) Effects of low light stress on dry matter accumulation and distribution of summer maize in 2021. S: Shading from flowering to maturity stage; CK: Normal light from flowering to maturity stage; PAA: The amount of assimilated dry matter after flowering; TBA: Dry matter translocation of pre-flowering; TFGR: Contribution rate of pre-flowering dry matter to grain; TRBA: Dry matter transport rate of pre-flowering; PAGR: Contribution rate of dry matter assimilation to grain after flowering. Different lowercase letters in the same column indicate significant difference at P < 0. 05 by LSD test.

### Gas exchange parameters in ear leaf

The photosynthetic rate of summer maize was significantly reduced under low light stress, and the change trend was consistent in the two years (Table 3). The Ci, Gs, Pn and E of S treatment were 22.8 %, 38.0 %, 19.5 % and 18.5 % lower than those of CK on average.

**Table 3.** Effects of low-light stress on gas exchange parameters in ear leaves of summer maize. Different lowercase letters in the same column indicate significant difference at P < 0. 05 by LSD test.

| Year | Treatment | Ci<br>($\mu\text{mol mol}^{-1}$) | Gs<br>($\text{mmol m}^{-2} \text{s}^{-1}$) | P<sub>n</sub><br>($\mu\text{mol m}^{-2} \text{s}^{-1}$) | E<br>($\text{mmol m}^{-2} \text{s}^{-1}$) |
| --- | --- | --- | --- | --- | --- |
| 2020 | CK | 125.0a | 339.7a | 34.5a | 7.4a |
|  | S | 93.5b | 210.5b | 27.1b | 5.9b |
| 2021 | CK | 111.1a | 406.0a | 36.0a | 6.6a |
|  | S | 88.5b | 252.0b | 29.7b | 5.5b |

S: Shading from flowering to maturity stage; CK: Normal light from flowering to maturity stage. Ci: Inter cellular carbon dioxide concentration; Gs: Stomatal conductance; P<sub>n</sub>: Net photosynthetic rate; E: Transpiration rate. Different lowercase letters in the same column indicate significant difference at $P < 0.05$ by LSD test.

### Observation of Kranz anatomy and determination of plasmodesmata densities

After low light stress, the number and area of Kranz anatomy in the rank-2 intermediate veins of the ear leaf were decreased. Compared with CK, the spacing of two Kranz anatomy in S treatment increased by 19.0 %, the average area of Kranz anatomy decreased by 9.9 %, and the density of Kranz anatomy per unit length decreased by 19.4 % (Figure 3).

**Figure 3.**
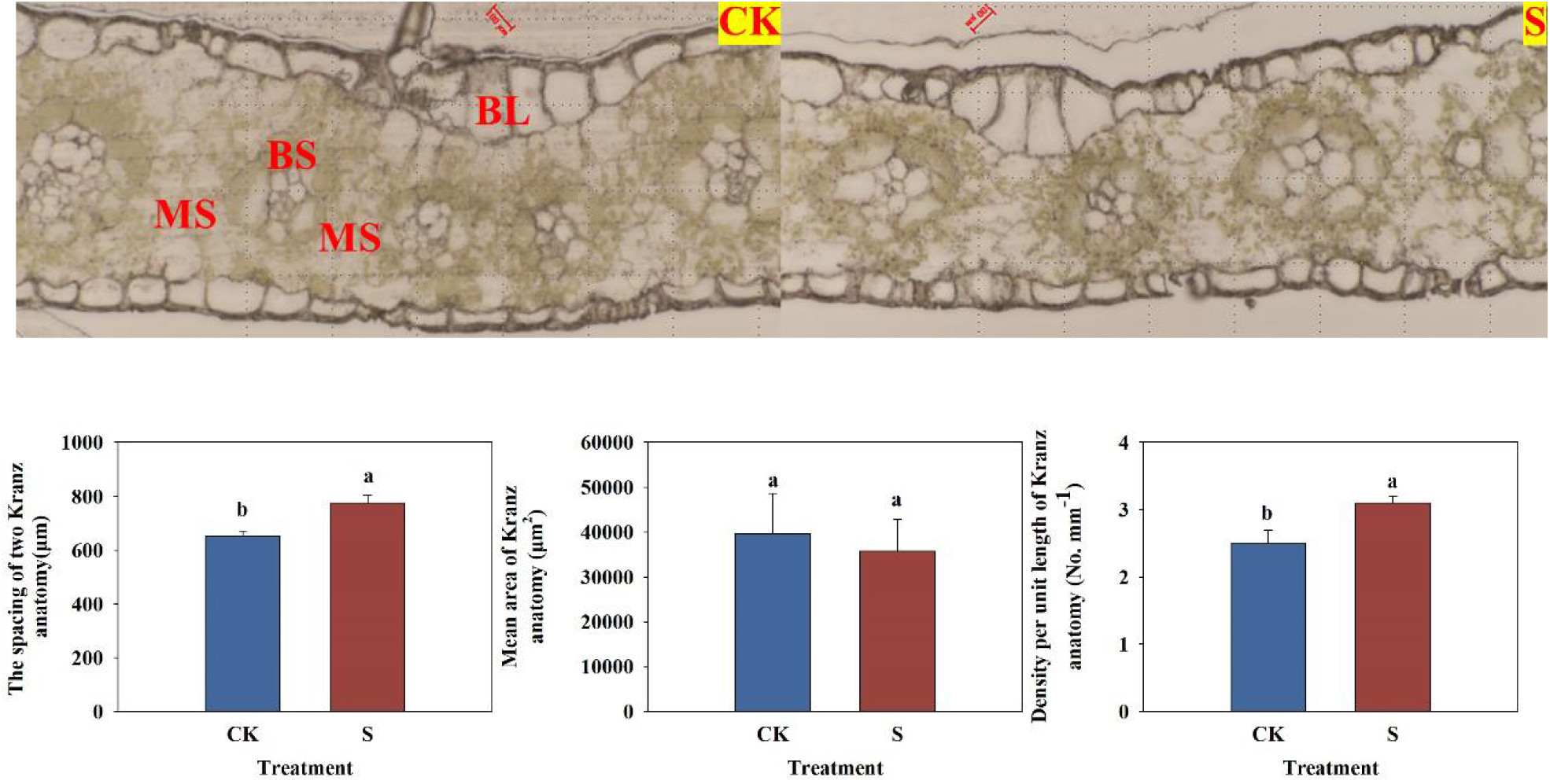
Effects of low-light stress on the number and area of Kranz anatomy in rank-2 intermediate veins of summer maize. S: Shading from flowering to maturity stage; CK: Normal light from flowering to maturity stage. BL: Bulliform cell; MS: Mesophyll cell; BS: Bundle sheath cell. The scale bar is 100 μm. Different lowercase letters in the same column indicate significant difference at P < 0. 05 by LSD test.

There were many chloroplasts and ordered arrangement in vascular bundle sheath cells treated with CK, and the number of starch granules was increased. There were many mitochondria in PP; the companion cells had dense cytoplasm, rich mitochondria and endoplasmic reticulum. The number of chloroplasts and starch granules in vascular bundle sheath cells treated with S was decreased; few mitochondria in PP; the companion cells were obviously vacuolated, containing a small amount of mitochondria and endoplasmic reticulum. There was no significant difference in the density of plasmodesmata between MS, BS and PP (Figure 4, 5).

**Figure 4.**
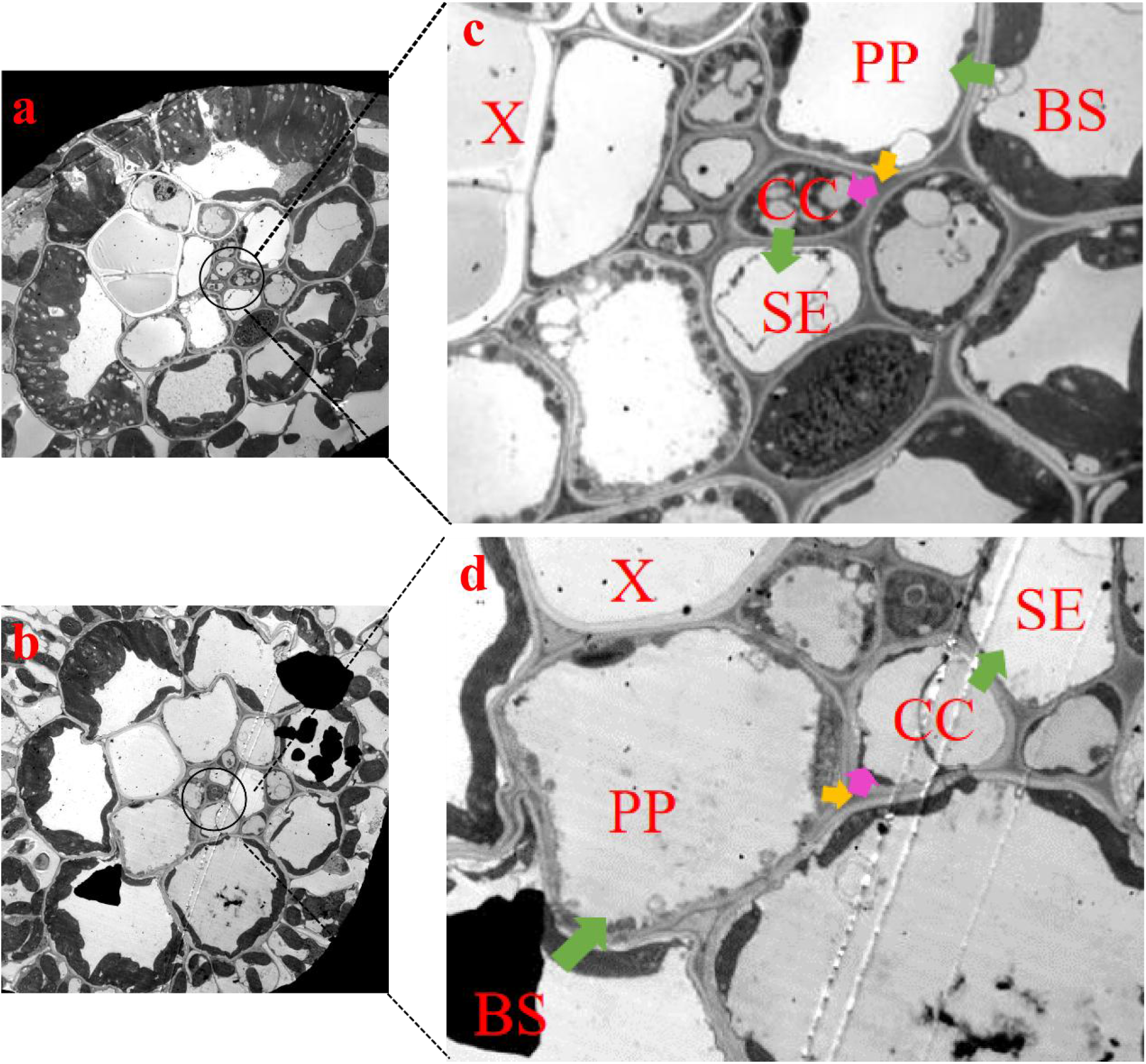
Effects of low-light stress on ultrastructure of summer maize leaves (2500×). a, c: CK; b, d: S. S: Shading from flowering to maturity stage; CK: Normal light from flowering to maturity stage. BS: Bundle sheath cell; PP: Phloem parenchyma cell; CC: Companion cell; SE: Sieve element; X: Xylem.

**Figure 5.**
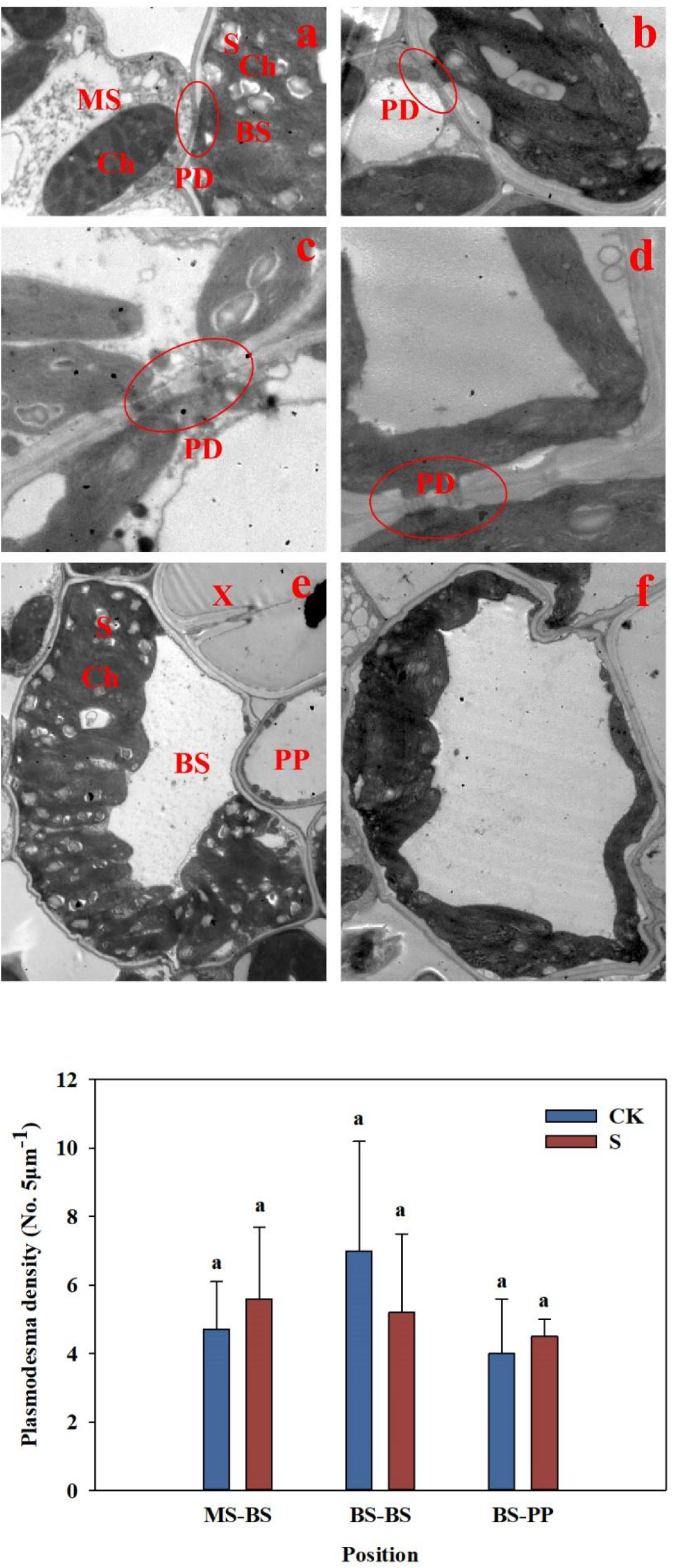
Effects of low-light stress on the number of plasmodesmata in leaves of summer maize. a, c, e: Denotes the plasmodesmata between MS and BS, the plasmodesmata between BS and BS, and the number of starch grains in BS of CK; b, d, f: Denotes the plasmodesmata between MS and BS, the plasmodesmata between BS and BS, and the number of starch grains of S. S: Shading from flowering to maturity stage; CK: Normal light from flowering to maturity stage. BS: Bundle sheath cell; MS: Mesophyll cell; PD: Plasmodesmata; PP: Phloem parenchyma cell; S: Starch grain; Ch: Chloroplast; X: Xylem. Different lowercase letters in the same column indicate significant difference at P < 0. 05 by LSD test.

### Sucrose transporter expression

Compared with CK, S treatment increased the relative expression of *SWEET13b* in leaves, while there was no significant difference in the relative expression of *SUT1, SWEET13a* and *SWEET13c* (Figure 6).

**Figure 6.**
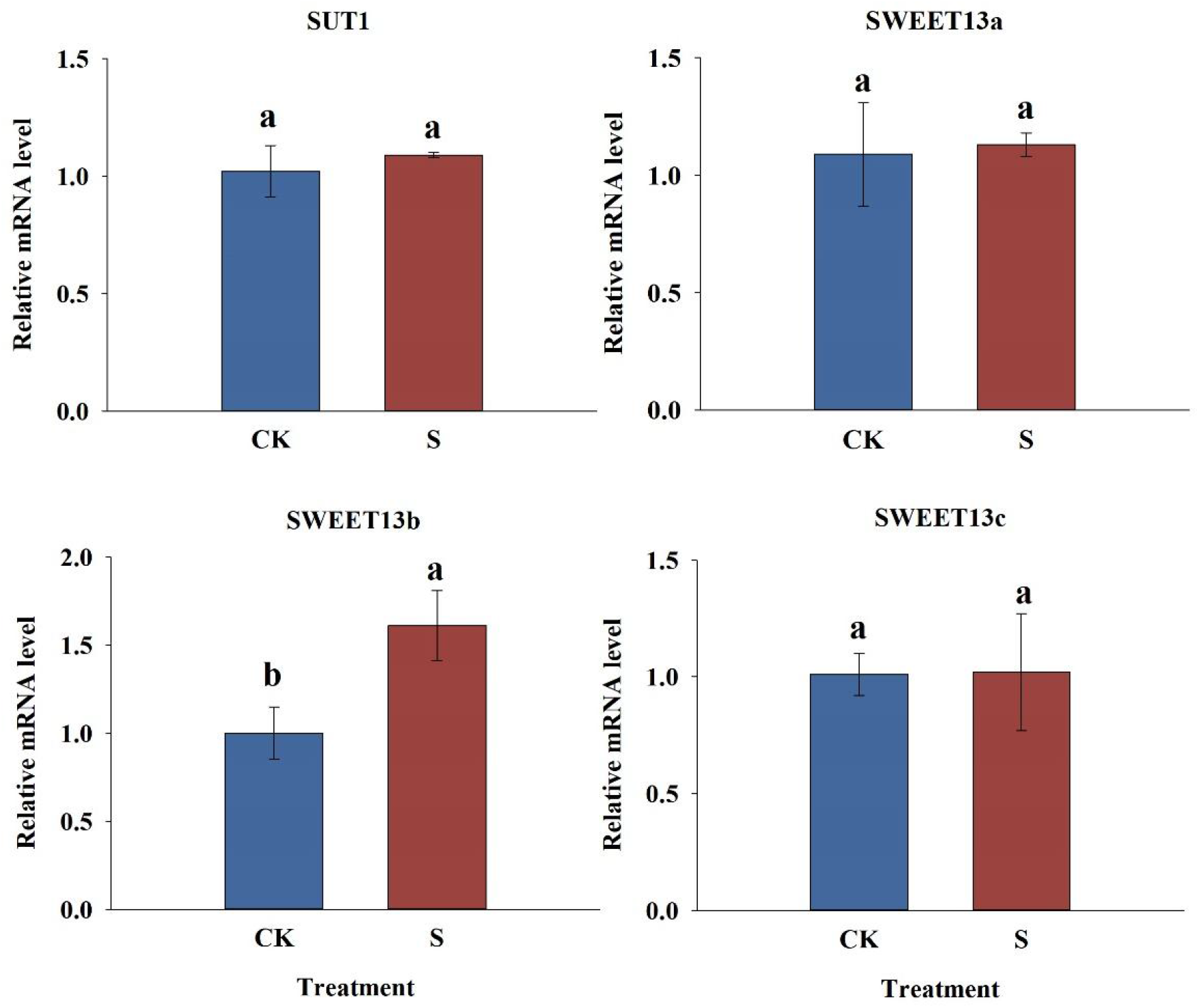
Effects of low-light stress on relative mRNA levels of sucrose transporter in summer maize leaves. S: Shading from flowering to maturity stage; CK: Normal light from flowering to maturity stage. The names of sucrose transporter are indicated at the top. Different lowercase letters in the same column indicate significant difference at P < 0. 05 by LSD test.

### Number and area of vascular bundles

The number of vascular bundles and the area of small vascular bundles were significantly reduced under low light stress. The area of large vascular bundles in shank was significantly reduced under low light stress, and there was no significant difference in the number of vascular bundles. The number of large and small vascular bundles in spike node of S treatment was 25.0% and 19.0% lower than that of CK; the total area, xylem area and phloem area of large vascular bundle were 1.8%, 8.2% and -0.4% lower than those of CK; small vascular bundle area decreased by 32.3% compared with CK. The number of large and small vascular bundles in shank of S treatment was 5.6% and 6.0% lower than that of CK. The total area, xylem area and phloem area of large vascular bundle were 26.7%, 10.3% and 32.5% lower than those of CK; the area of small vascular bundle was 1.0% lower than that of CK (Figure 7).

**Figure 7.**
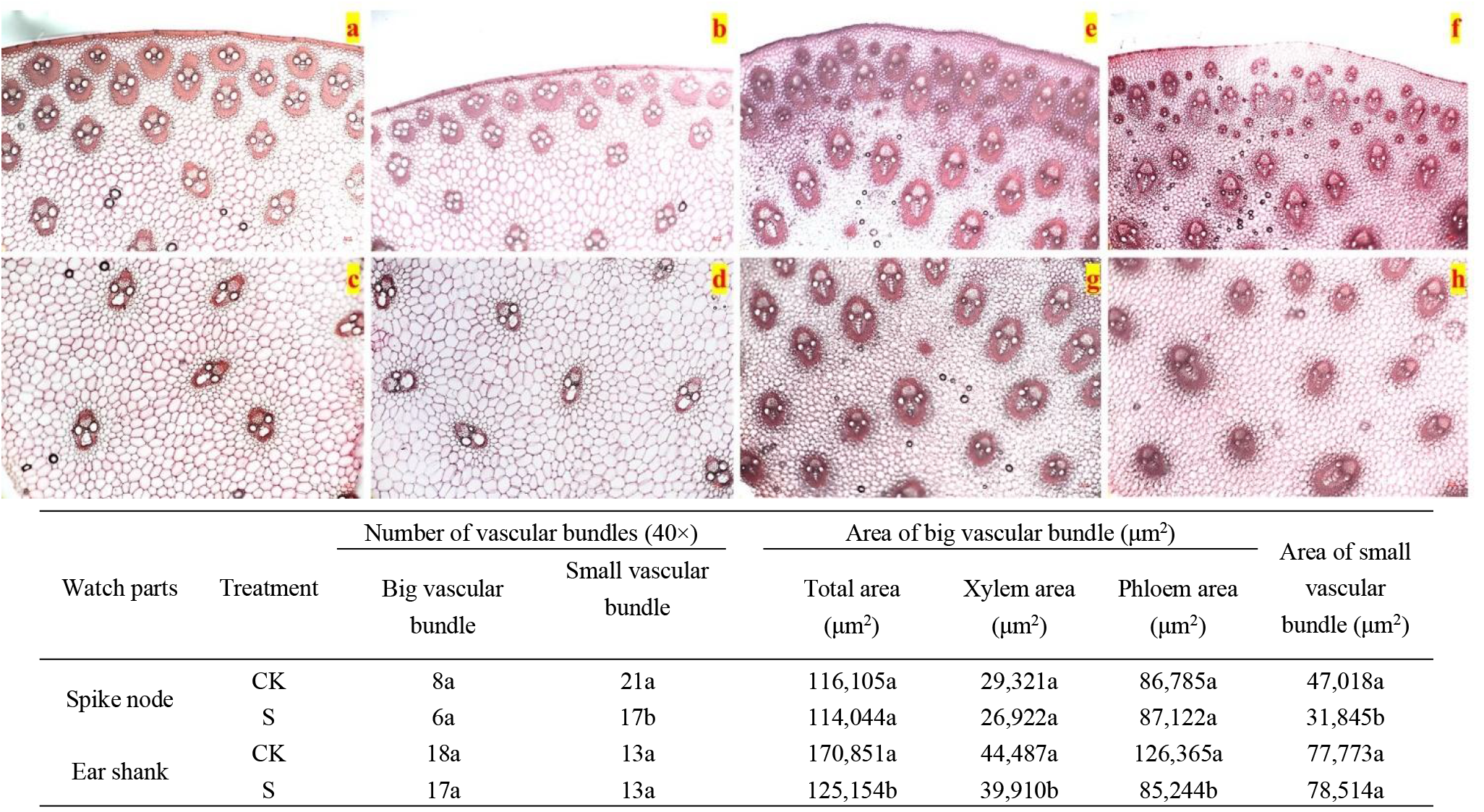
Effects of low-light stress on the structure of small vascular bundle and central vascular bundle of spike node and ear shank in summer maize (40×). a, c: Denotes the structure of small vascular bundle and central vascular bundle of spike node of CK. b, d: Denotes the structure of small vascular bundle and central vascular bundle of spike node of S. e, g: Denotes the structure of small vascular bundle and central vascular bundle of ear shank of CK. f, h: Denotes the structure of small vascular bundle and central vascular bundle of ear shank of S. S: Shading from flowering to maturity stage; CK: Normal light from flowering to maturity stage. Different lowercase letters in the same column indicate significant difference at P < 0. 05 by LSD test.

### Grain filling parameters

Maximum grain filling rate was significantly reduced and active grain filling period was shortened under low light stress. Tmax, Wmax, Gmax, Gmean and D of S treatment were -6.6%, 32.6%, 28.4%, 0.1% and 4.4% lower than CK (Figure 1).

### Concentrations of sucrose and starch

The contents of sucrose and starch in ear leaves and grains of summer maize was changed under low light stress. In 2021, the concentrations of sucrose, starch and the ratio of starch to sucrose (S / S) in ear leaves and grains of S treatment were 41.5%, 85.1%, -5.9% and 97.3%, 509.9%, 19.7% lower than those of CK, respectively. From the sucrose concentration gradient between source and sink, the sucrose concentration in ear leaves of S treatment was lower than that of CK, while the sucrose concentration in grains of S treatment was higher than that of CK (Figure 8).

**Figure 8.**
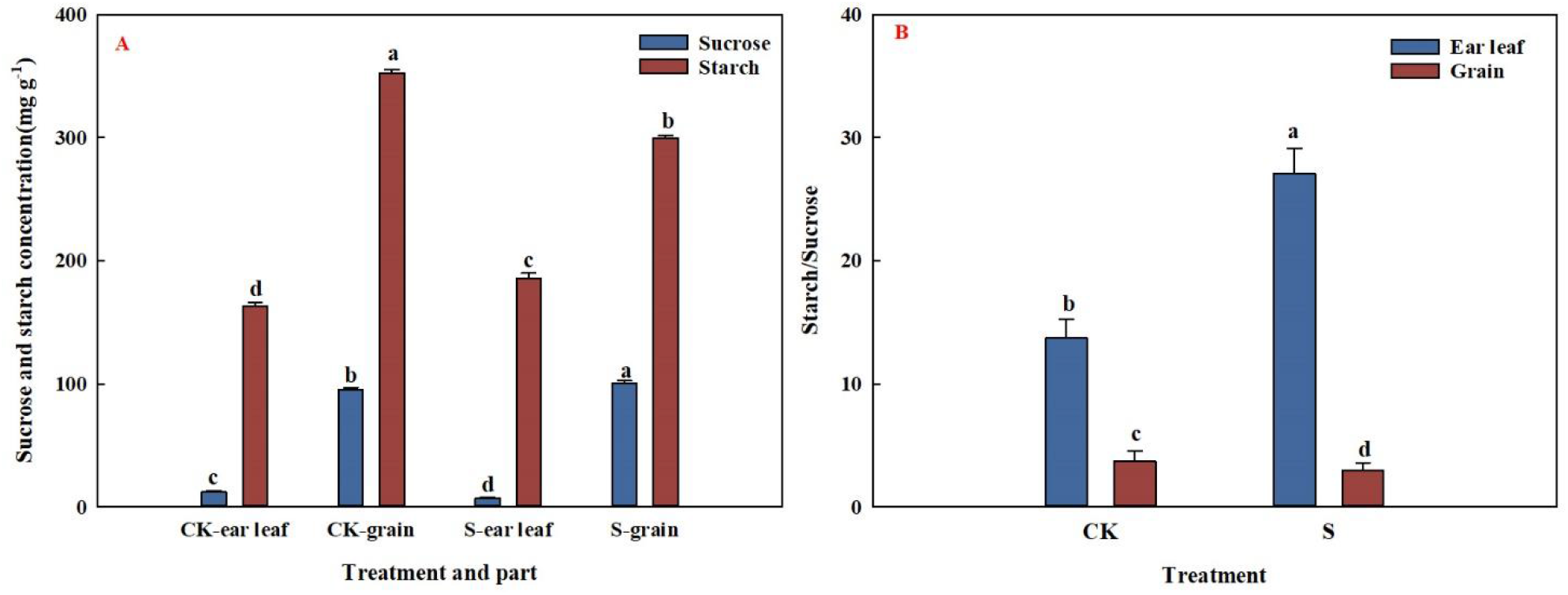
Effects of low-light stress on sucrose and starch contents in ear leaf and grain of summer maize. (A) Effects of low-light stress on sucrose and starch concentration in ear leaf and grain of summer maize. (B) Effects of low-light stress on the ratio of sucrose and starch in ear leaf and grain of summer maize. S: Shading from flowering to maturity stage; CK: Normal light from flowering to maturity stage. Different lowercase letters in the same column indicate significant difference at P < 0. 05 by LSD test.

### Sucrose-starch metabolic enzyme activity

Low light stress significantly reduced Rubisco, PEPC, NADP-ME, SPS enzyme activities in ear leaves and SUS, AGPase enzyme activities, fructose content in grains. The activities of Rubisco, PEPC, NADP-ME, SPS and AGPase in ear leaves of S treatment were decreased by 25.2 %, 47.8 %, 63.0 %, 35.8 % and 3.7 % compared with CK. Compared with CK, the activities of SUS and AGPase and fructose content in grains of S treatment decreased by 15.5 %, 18.2 % and 10.46 %, respectively, and there was no significant difference in CWI activity (Figure 9).

**Figure 9.**
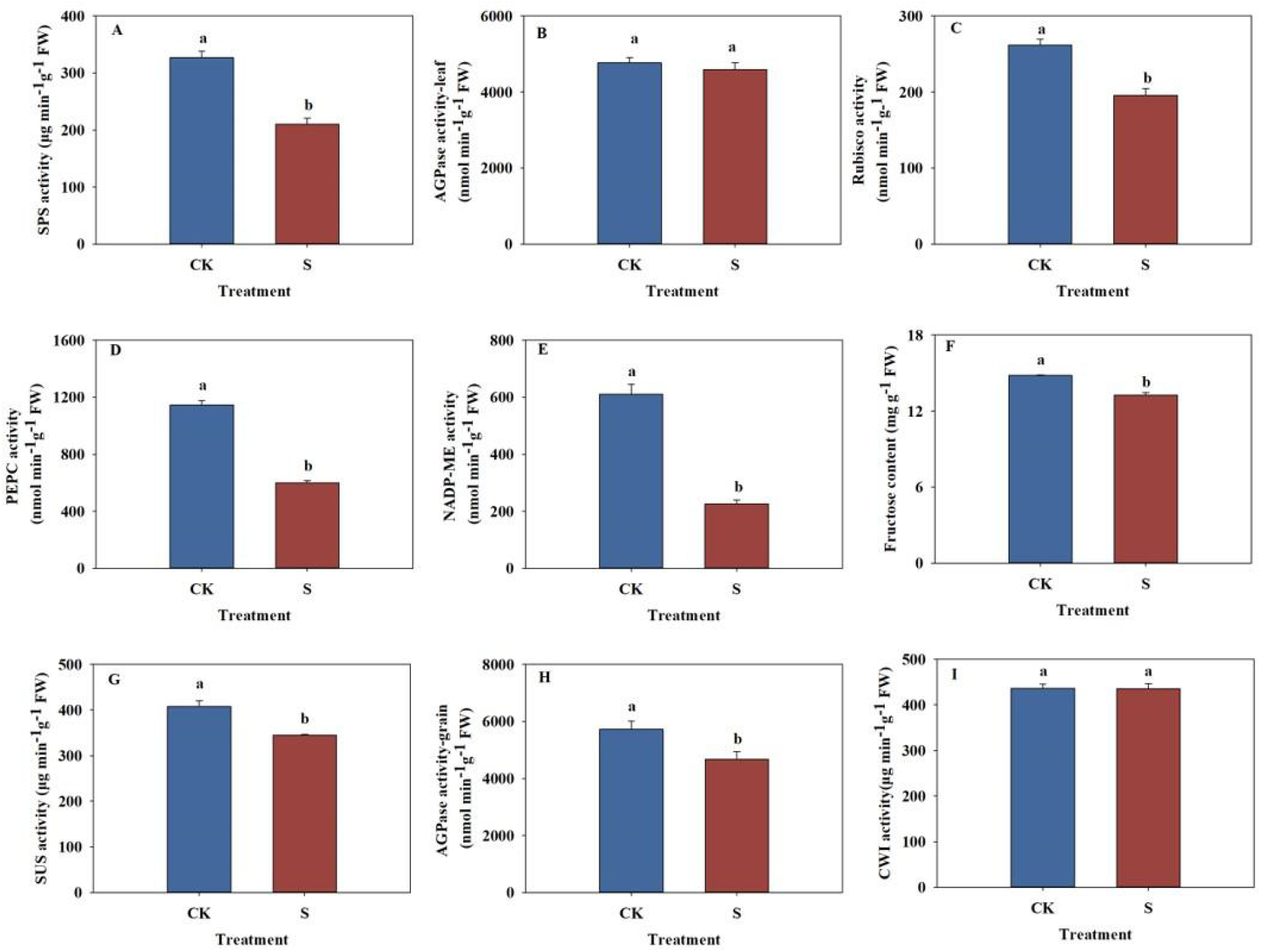
Effects of low-light stress on sucrose-starch metabolic enzyme activities in leaves and grains of summer maize. (A) Effects of low-light stress on SPS activities in leaves of summer maize. (B) Effects of low-light stress on AGPase activities in leaves of summer maize. (C) Effects of low-light stress on Rubisco activities in leaves of summer maize. (D) Effects of low-light stress on PEPC activities in leaves of summer maize. (E) Effects of low-light stress on NADP-ME activities in leaves of summer maize. (F) Effects of low-light stress on fructose content in grains of summer maize. (G) Effects of low-light stress on SUS activities in grains of summer maize. (H) Effects of low-light stress on AGPase activities in grains of summer maize. (I) Effects of low-light stress on CWI activities in grains of summer maize. SPS: Sucrose phosphate synthase; AGPase: Adenosine diphosphate glucose pyrophosphorylase; Rubisco: Ribulose-1,5-bisphosphate carboxylase/oxygenase; PEPC: Phosphoenolpyruvate carboxylase; NADP-ME: NADP-dependent malic enzyme; SUS: Sucrose synthase; CWI: Cell wall invertase; S: Shading from flowering to maturity stage; CK: Normal light from flowering to maturity stage. Different lowercase letters in the same column indicate significant difference at P < 0. 05 by LSD test.

## Discussion

### Low light changed the ability of leaves to synthesize sucrose and transport sucrose to grain

The main source of grain yield was photoassimilates formed after silking (Shi *et al*., 2015). The synthesis and activation of Rubisco, PEPC and NADP-ME were induced by light. This study showed that under a low light environment, the activities of NADP-ME, PEPC and Rubisco in leaves was decreased, and the Ci was decreased, resulting in the decrease of photosynthetic rate and the decrease of assimilates. The results were consistent with Zhang et al. (Zhang *et al*., 2007, 2008). The triose phosphate produced during the Calvin cycle was transported to the cytoplasm by the Triose phosphate / phosphate translocator (TPT), which synthesized sucrose by SPS and then transported to the grain. The results showed that the activity of AGPase remained unchanged and the activity of SPS was decreased after shading, resulting in the decrease of sucrose content, the increase of starch content and the increase of the ratio of starch to sucrose. The results were consistent with Liang et al. (Liang *et al*., 2020). The activity of TPT was affected by light (Wang *et al*., 2002), and high concentration of inorganic phosphate (Pi) inhibited the activities of AGPase and SPS. Ning et al. have shown that nitrogen deficiency in leaves led to low activity of TPT, resulting in reduced triose phosphate transported out of chloroplasts and reduced Pi transported into chloroplasts (Ning *et al*., 2018). Therefore, it was speculated that under low light conditions, the expression level of TPT in leaves was low, resulting in a decrease in triose phosphate transported to cytoplasm, and an increase in the proportion of RuBP regeneration and starch synthesis. In addition, the triose phosphate in cytoplasm decreased, the stimulation on SPS activity decreased, and the synthesis of sucrose was blocked, resulting in an increase in the ratio of starch to sucrose. However, it is difficult to determine the output rate of triose phosphate at present, and the above conjecture needs further verification in the future. Bundle sheath cells in Rank-2 intermediate veins were closely linked to mesophyll cells, mainly responsible for sucrose loading into phloem. However, low light down-regulated the expression of enzymes related to glycolysis (Gao *et al*., 2020), inhibited glycolysis metabolism and lacked energy, leading to premature senescence of leaves (Huang *et al*., 2020; Wu *et al*., 2021), in which bundle sheath cells senescence faster than mesophyll cells (Wu *et al*., 2021). In this study, we also found that under low light stress, the number and area of Kranz anatomy was reduced, the number of chloroplasts in bundle sheath cells was decreased, and the number of mitochondria in PP was decreased, accompanied by obvious vacuoles, which was consistent with the symptoms of premature senescence. In addition, energy consumption was required for sucrose efflux and entry the CC-SE complex during active apoplasmic loading of summer maize (Bezrutczyk *et al*., 2021). Under low light conditions, the number of mitochondria in PP cells and companion cells decreased, and the energy supply was insufficient. It was difficult for sucrose to move to SE, and the output power was small. Chen et al. have shown that SWEET13 in Arabidopsis protected the normal efflux of sucrose under low light conditions (Chen *et al*., 2011). Interestingly, this study also found that the frequency of plasmodesmata between MS-BS-PP and the expression of transporters responsible for efflux and absorption of sucrose were not affected by low light, which might be a manifestation of summer maize adapting to low light. Mathan et al. (2021) quantitatively analyzed the sucrose loading capacity in the phloem of detached rice leaves by detecting the isotope content. In the future, more convenient and accurate methods are needed to determine the sucrose concentration and sucrose loading rate in the phloem of maize leaves under low light conditions to explain the mechanism of source-sink relationship change. In conclusion, the synthesis ability of sucrose in leaves and the export sucrose to grains ability of leaves were reduced under low light.

### In low light conditions, stem dry matter transport was increased, but due to this, long-distance transport of light contract compounds was limited

Part of the dry matter required for grain filling came from the dry matter transported by stems, and the other part came from the dry matter synthesized by leaves. This study showed that under low light conditions, when the dry matter produced by leaves could not meet the grain filling, the dry matter accumulation before flowering in vegetative organs was increased, and its contribution rate to grain was greater than that of photosynthetic assimilation after flowering to grain. This was consistent with the results of Wang et al. (2020) and Yang et al. (2021). Our previous research group also obtained (Gao *et al*., 2017) that the contribution rate of stem-transported dry matter to grain accounted for 86 % of the contribution rate of pre-flowering dry matter transport to grain yield, and the spike node and its upper and lower nodes were the most transported. Although dry matter transport in stem was increased, this adaptation mechanism could not compensate for the loss of yield and had a negative impact on long-distance transport of photosynthetic assimilates. Previous studies have found that the number and area of vascular bundles were closely related to the non-structural carbohydrate transport, seed setting rate, harvest index and yield of stem and sheath, and small vascular bundles contributed more to yield due to the high density of plasmodesmata in phloem (Li *et al*., 2019). In this experiment, we found that the number and area of small and medium-sized vascular bundles in spike node and the area of large vascular bundles in shank were reduced under low light, which resulted in the long-distance transport of sucrose in stem was limited, the transport rate was decreased, the “flow” was not smooth, affecting the grain filling rate and filling time.

### Low light altered the utilize sucrose ability of grains

After long-distance transportation of sucrose to grains, on the one hand, it was hydrolyzed by CWI and entered cells under the action of transporters. On the other hand, it was directly transported into cells by sucrose transporters (Zhang *et al*., 2018). After sucrose entered the grain, starch was synthesized through a series of enzymes. The results showed that CWI activity was remained unchanged, but the activities of SUS and AGPase were decreased, resulting in the decrease of fructose and starch content. The results were consistent with research results of Zhang et al. (2008). The process of leaf synthesis, sucrose loading and sucrose utilization in grains was affected under low light, resulting in relatively higher sucrose concentration in grains than in leaves, forming a “leaf low” - “grain high” sugar concentration gradient, resulting in the opposite hydrostatic pressure, and then feedback inhibition of sucrose output in leaves, reducing sucrose loading and transportation rate (Figure 10). In summary, the direction of dry matter distribution was changed under low light, and the insufficient supply of dry matter and low photoassimilation ability led to insufficient grain plumpness, decreased grain number per ear and grain weight, and significantly reduced grain yield. Among them, the decrease of leaf photosynthetic rate was the main reason for yield reduction. The key to solving this problem in the future is to improve the photosynthetic capacity of leaves by screening shade-tolerant varieties or cultivation measures (such as increasing nitrogen fertilizer, removing top leaves, spraying growth regulators and foliar fertilizer), ensure the supply of sucrose in grains, alleviate the competition between leaves and grains for limited sucrose, and achieve stable and high yield of summer maize.

**Figure 10.**
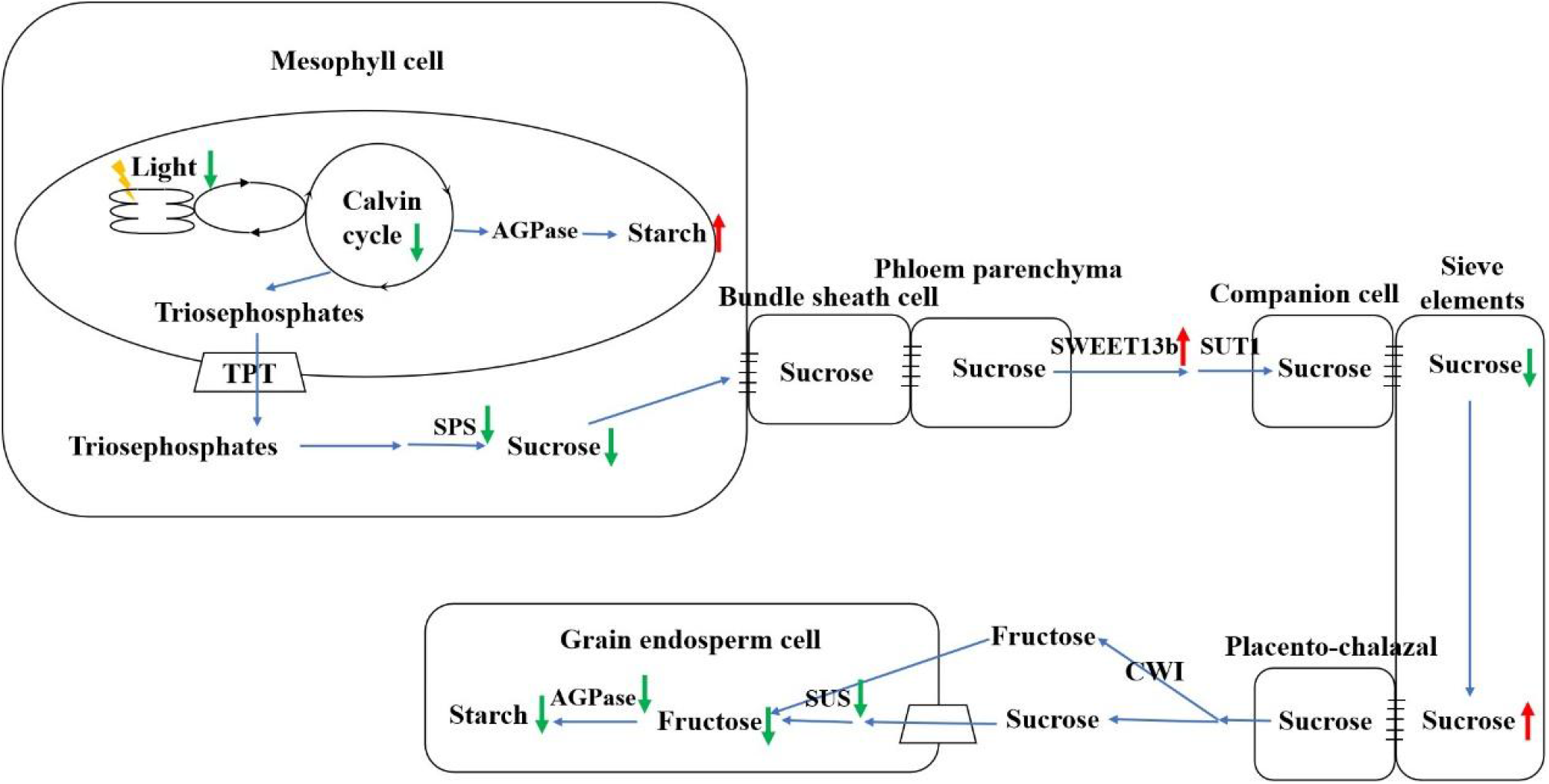
Effects of low-light stress on sucrose transport efficiency of summer maize. TPT: Triose phosphate/phosphate translocator; SPS: Sucrose phosphatase; CWI: Cell wall invertase; SUS: Sucrose synthase; AGPase: Adenosine diphosphate glucose pyrophosphorylase.

## Conclusions

Due to insufficient light, the dry matter production after flowering of summer maize leaves was seriously decreased, the number and area of Kranz anatomy were reduced, and the phloem cells were vacuolated, resulting in the decrease of sucrose synthesis in leaves and the export sucrose to grains ability of leaves. Stem dry matter transport was increased before flowering in order to meet the needs of grain filling, but this adaptation mechanism could not compensate for the loss of yield, and also led to the decrease of the number and area of stem vascular bundles, the obstruction of dry matter transport and the decrease of grain filling rate. Yield was significantly decreased under low light stress by affecting photoassimilates synthesis, distribution of photoassimilates from leaf to ear, transportation of photoassimilates from stem to ear and utilization of photoassimilates in grain.

## Author Contribution

Jiwang Zhang and Zhichao Sun involved in to conceptualization. Zhichao Sun and Wenjie Geng involved in methodology. Baizhao Ren, Peng Liu and Bin Zhao involved in formal analysis. Zhichao Sun and Wenjie Geng involved in investigation. Jiwang Zhang involved in resources. Zhichao Sun and Wenjie Geng involved in data curation. Zhichao Sun and Jiwang Zhang writing—original draft preparation and writing—review and editing. All the authors have read and agreed to the published version of the manuscript.

## Data Availability

All data supporting the findings of this study are available within the paper and within its supplementary materials published online. Further inquiries can be directed to the corresponding author (Zhang Jiwang), upon request.

